# Optimized design of antisense oligomers for targeted rRNA depletion

**DOI:** 10.1101/2020.06.24.169102

**Authors:** Wesley A. Phelps, Anne E. Carlson, Miler T. Lee

**Affiliations:** Department of Biological Sciences, University of Pittsburgh, Pittsburgh PA 15260 USA

## Abstract

RNA sequencing (RNA-seq) has become a standard method for quantifying gene expression transcriptome-wide. Although RNA-seq is often paired with polyadenylate (poly(A)) selection to enrich for messenger RNA (mRNA), many applications require alternate approaches to counteract the high proportion of ribosomal RNA (rRNA) in total RNA. Recently, selective rRNA digestion, using RNaseH and antisense DNA oligomers that tile the entire length of target RNAs, has emerged as an alternative to commercial rRNA depletion kits. Here, we present a streamlined, more economical RNaseH-mediated rRNA depletion method with substantially lower up-front costs, using shorter antisense oligos only sparsely tiled along the target RNA, in a digestion reaction of only 5 minutes. We introduce a novel Web tool, Oligo-ASST, that simplifies oligo design to target regions with optimal thermodynamic properties, and additionally can generate compact, common oligo pools that simultaneously target divergent RNAs, e.g. across different species. We demonstrate the efficacy of these strategies by designing oligo sets to deplete rRNA in *Xenopus laevis* and in zebrafish, which expresses two distinct versions of rRNAs during embryogenesis. The resulting RNA-seq libraries reduce rRNA to <5% of aligned reads, on par with poly(A) selection, and also reveal expression of many non-adenylated RNA species. Oligo-ASST is freely available at https://mtleelab.pitt.edu/oligo to design antisense oligos for any taxon or to target any abundant RNA for depletion.

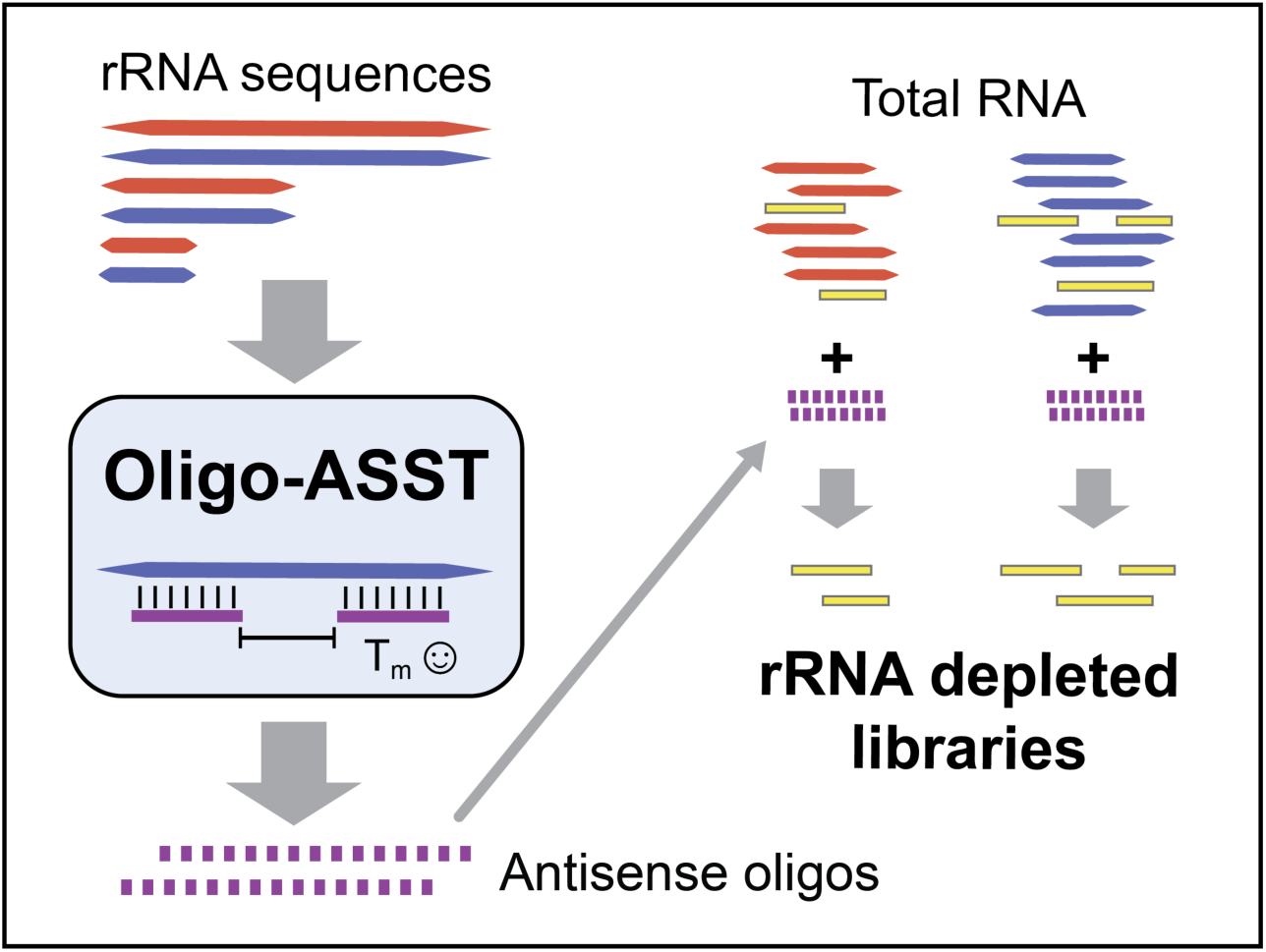

## INTRODUCTION

High-throughput RNA sequencing (RNA-seq) has become a widespread method for measuring gene expression transcriptome-wide[1]. Most RNA-seq studies focus on messenger RNA (mRNA); however, the vast majority of total RNA (>80%) [2, 3] is ribosomal RNA (rRNA). Therefore, RNA-seq is commonly paired with methods to reduce the amount of rRNA included in sequencing libraries, to maximize the proportion of sequencing reads derived from genes of interest.

An effective, widely used strategy for enriching mRNA is polyadenylate (poly(A)) selection (poly(A)+)[1]. In eukaryotes, most mRNAs encode 3’ poly(A) tails, which are used to select for and enrich mRNA pools using oligo(dT)-based methods[1, 4]. However, many applications cannot take advantage of this approach, notably transcriptomics in prokaryotes, whose mRNA largely lack poly(A) tails[3], but also many eukaryotic contexts as well. Methods that aim to quantify message fragments separated from poly(A) tails, such as RNA-seq on degraded RNAs[5, 6], cap analysis gene expression (CAGE)[7] and ribosome profiling[8], require alternate rRNA depletion strategies. Some RNAs of interest, such as nascent pre-mRNA, some histone mRNA and many non-coding RNAs (ncRNAs), do not encode poly(A) tails[9], thus their expression levels are underrepresented in poly(A)+ RNA-seq libraries. Finally, since poly(A) tail length is variable, it is a challenge to distinguish changes in poly(A) status, e.g. due to the activity of deadenylases, from changes in RNA molecule number using poly(A)+ RNA-seq[10]. Indeed, in animals such as *Xenopus* and zebrafish, the maternal mRNA contribution to the egg is largely deadenylated[11], thus poly(A) selection is not well suited to accurately measure the transcriptome in the early embryo[12-15].

An alternative to enrich for mRNAs in libraries is exon capture, in which samples are hybridized to oligo probes designed against the transcriptome of interest prior to sequencing [16]. While particularly effective in clinical settings [17], commercial probe sets are costly and available for only select, well-annotated transcriptomes, limiting the feasibility of this method for many RNA-seq applications. Thus, methods to exclude unwanted RNA species have also been widely used, including electrophoretic size selection[18], digestion of highly abundant species in cDNA libraries using duplex-specific nucleases [19, 20], and targeted rRNA depletion[21]. In the latter approach, DNA oligos complementary to rRNA facilitate their removal prior to library construction. These oligos can be biotinylated for magnetic bead affinity purification [6, 22, 23], commercialized for some taxa in the Ribo-Minus and the now discontinued Ribo-Zero Gold kits.

Antisense oligos can be used in conjunction with RNaseH to digest DNA-rRNA hybrids[5, 24]. Several studies in both mammals and bacteria have shown that RNaseH-mediated rRNA depletion is efficient, resulting in sequencing libraries with minimal rRNA derived reads [5, 23-27]. Commercial solutions have emerged for select taxa; however, the ease of this method allows it in principle to be readily adapted to any taxon, with the primary challenge being the design and acquisition of the 50nt oligos that tile the specific rRNA sequences encoded in its transcriptome. Although rRNA sequences are generally well conserved between species relative to other genes, nucleotide differences and variable regions at even modest evolutionary distances[28] pose a challenge for reusing oligos designed for one taxon to effectively perform rRNA depletion in another.

Here, we present an optimized strategy for RNaseH-mediated rRNA depletion suitable for RNA-seq library construction that reduces up-front oligo costs by as much as 81%. Using *Xenopus laevis* and zebrafish (*Danio rerio*) embryos as test cases, we demonstrate that short (39-40nt) antisense DNA oligos sparsely tiled along rRNA, coupled with a 5-minute digestion, effectively produces RNA-seq libraries with <5% rRNA-derived reads, on par with poly(A) selection. We show that divergent rRNAs can be simultaneously digested with partially overlapping oligo pools that target regions of high sequence similarity, facilitating the design of flexible, cross-taxon reagents for rRNA depletion. Finally, we introduce a web tool, Oligo-ASST, that simplifies oligo design, allowing this approach to be easily adapted to any taxon or to target any other abundant RNAs for depletion.

## MATERIALS AND METHODS

### Animal husbandry

All animal procedures were conducted under the supervision and approval of the Institutional Animal Care and Use Committee at the University of Pittsburgh. *X. laevis* adults (NASCO NXR_0.0031) were housed in a recirculating aquatic system (Aquaneering) at 18°C with a 12/12 hour light/dark cycle. Frogs were fed twice weekly with Frog Brittle (NASCO #SA05960(LM)M). *Danio rerio* (zebrafish) were housed in a recirculating aquatic system (Aquaneering) at 27°C with a 14/10 hour light/dark cycle and fed freshly hatched *Artemia spp.* nauplii twice daily, supplemented with TetraMin Tropical Flakes and dried krill.

### Sample collection

To obtain *Xenopus laevis* embryos, sexually mature females were injected with 1000 IU human chorionic gonadotropin into their dorsal lymph sac and incubated overnight at 16°C. In the morning, females were moved to room temperature where they laid eggs within an hour of being moved. Sexually mature males were euthanized by 30 minute submersion in 3.6 g/L tricaine-S (MS-222), pH=7.4, and testes were dissected. Cleaned testes were stored up to a week in L-15 medium at 4°C. Eggs were collected and artificially inseminated in MR/3 (33 mM NaCl, 0.6 mM KCl, 0.67 mM CaCl_2_, 0.33 mM MgCl_2_, 1.67 mM HEPES, pH 7.8)[29]. Zygotes were de-jellied[30] in MR/3 pH=8.5, with 0.3% β-mercaptoethanol with gentle manual agitation, neutralized with MR/3 pH=6.5, washed twice with MR/3 and incubated in MR/3 at 23°C until desired developmental stage.

Zebrafish embryos were obtained from natural mating of TUAB strain fish 6-12 month old. Mating pairs were selected randomly from a pool 24 males and 24 females >=1 month since last breeding. Zebrafish were isolated in mating pairs overnight at room temperature in divided tanks. Dividers were removed the following morning, and eggs were collected in egg water (60 μg/ml Ocean salt in RO water) and incubated at 28.5°C until the desired developmental stage.

To obtain fin clips, adult zebrafish were anesthetized in 500mg/L MS-222 in system water for 2-5 minutes until gills stopped moving, then one lobe of the caudal fin was clipped. Fish were transferred to fresh system water for recovery.

### Total RNA extraction

For *X. laevis*, 2 embryos were pooled for RNA extraction; for *D. rerio*, 20 embryos or 10 fin clips were pooled. Samples were snap frozen in a 1.5μl tube and homogenized with a pestle in 500μl of TRIzol Reagent (Invitrogen #15596026) followed by 100μl of chloroform. Tubes were centrifuged at 18,000 x g at 4°C for 15 minutes, the aqueous phase was transferred to a fresh tube with 340 μl of isopropanol and 1 μl of GlycoBlue (Invitrogen #AM9515), then precipitated at -80°C for 1 hour. Precipitated RNA was washed with cold 75% ethanol and resuspended in 50μl of nuclease-free water. Concentration was determined by NanoDrop. RNA was stored at -80°C until use.

### Antisense oligo design

The oligo tiling program is written in Python3. Each tiled oligo is defined by the start and end position of the complementary region in the target sequence (e.g., an rRNA). The algorithm assigns oligo positions left to right in a greedy fashion, such that each oligo is the maximum distance from the previous placed oligo while satisfying the parameter constraints – by default, melting temperature (Tm) between 70-80°C, length between 39-40 nucleotides (nts), and maximum untiled region ≤30 nts. If no oligo exists that satisfies these constraints, the oligo with closest Tm to the allowable range is retained. The maximum untiled region is iteratively adjusted to take into account the remaining sequence length. Melting temperature is calculated using the nearest-neighbor method[31, 32] with RNA-DNA parameters[33] assuming a Na^+^ concentration of 200mM and a conservatively low oligo concentration of 50nM (which will yield Tms 1-2°C lower compared to the highest oligo concentration we use, 400nM). Once the entire target sequence is tiled, a second refinement phase adjusts each oligo position within the window defined by the upstream and downstream gaps, to yield maximized distances from upstream and downstream oligos within the optimal Tm range.

To find shared oligo pools between 2 or more unaligned target sequences, oligo tiling proceeds as above for the first sequence. For each subsequent sequence, oligos from the first set with exact complementary matches are selected, then the remaining untiled regions are subjected to the tiling procedure as above. To find shared oligo pools between aligned target sequences, a consensus sequence from the alignment is used for the first round of oligo tiling to generate the candidate common oligos for subsequent rounds of tiling for each individual sequence. If wildcards bases are allowed, the consensus sequence will incorporate IUPAC wildcard bases. Wildcard-containing oligos are retained if the number of possible target sequences does not exceed the threshold specified by the user (e.g., an oligo with two wildcard positions, R and Y, would target four different sequences encoding all combinations of A/G and C/T at the complementary positions).

Oligos for *X. laevis* rRNA were designed individually for 28S (X02995.1:3836-7917), 18S (X02995.1:1030-2854), 5.8S (X02995.1:3412-3573), 16S (M10217.1:3093-4723), and 12S (M10217.1:2205-3023). Aligned consensus oligos were designed for the maternal and somatic 5S (maternal: M10635:352-471, somatic: J01009.1:607-726)[34]. For zebrafish, aligned consensus oligos were designed for maternal and somatic 28S (chr4:77556054-77560323(-) and chr5:820029-824137(-) respectively), maternal and somatic 18S (chr4:77561203-77563141(-) and chr5:824921-826807(-) respectively), maternal and somatic 5.8S (chr4:77560653-77560810(-) and chr5:824488-824644(-) respectively), and maternal and somatic 5S (chr4:41890222-41890340(-) and chr18:30048558-30048676(-) respectively), according to previous annotations[35, 36]. Individual oligos sets were designed for 16S (chrM:1020-1971(+)) and 12S (chrM:2043-3725(+)). All coordinates are from the GRCz11 genome build.

Oligos were ordered from Thermo Fisher as individual dry, desalted tubes at 25nM scale. At the time of writing, value oligo pricing (≥25 oligos with length ≤40 nts) was $4.64 per oligo, thus a full *X. laevis* set (137 oligos) would cost ∼$636. In contrast, standard 50mer oligos are $19 each (without institutional discount), thus the 176 oligos required for full tiling would total $3344. With an institutional discount, this would likely still be >$1400.

### RNaseH-mediated depletion

Individual dry oligos were resuspended to 1000 μM. For *X. laevis*, a 10X working stock for nuclear rRNA (28S, 18S, 5.8S, maternal and somatic 5S) was created by pooling 1μl of each of the 96 oligos and diluting to 4 μM per individual oligo (250 μL total volume, 384 μM total oligo concentration). At 1X concentration in 10μl, each oligo is at 400nM, which we estimate to be 10-fold in excess of its target in 1μg of total RNA: assuming 80% of total RNA is derived from 40S rRNA (28S, 18S, 5.8S in equimolar amounts) and 28S rRNA is 2x the length of 18S+5.8S, this corresponds to ∼530ng of the ∼4000-nt 28S rRNA, or ∼41nM in 10μl. A similar stock of 41 oligos targeting the less abundant mitochondrial rRNA (16S, 12S) was prepared at 1μM per individual oligo. For zebrafish, separate working stocks for maternal nuclear (112 oligos at 4μM per oligo), somatic nuclear (109 oligos at 4μM per oligo), and mitochondrial rRNA (42 oligos at 1μM per oligo) were similarly constructed. Maternal and somatic nuclear pools were then proportionally mixed according to developmental stage (1:0 for 2-cell, 1:1 for 28hpf, 0:1 for adult)[36]. Hybridization procedure was based on Adiconis et. al.[24] with slight modifications: 1μl of the nuclear pool (final concentration 0.4 μM per oligo) and 1μl of the mitochondrial pool (final concentration 0.1 μM per oligo) were combined with 1μg of total RNA (and optionally 150 ng of in vitro transcribed mCherry mRNA) in a 10μl buffered reaction volume (100mM Tris-HCl pH 7.4, 200mM NaCl, 10mM DTT), heated at 95°C for 2 minutes and cooled to 22°C at a rate of 0.1°C/s in a thermocycler. Next, 10U of thermostable RNaseH (NEB #M0523S) and 2μl of provided 10X RNaseH buffer were added and volume brought to 20μl with nuclease-free water. We achieved the best results with NEB thermostable RNaseH compared to other commercial RNaseH products. The reaction was incubated at either 45°C or 65°C for 5 or 30 minutes, then 5U of TURBO DNase (Invitrogen #AM2238) and 5μl of provided 10x buffer was added, volume brought to 50μl with nuclease-free water and incubated at 37°C for 30 minutes. Oligos were omitted from input control samples prior to heating and enzyme addition. For visualization, 12.5 μl of each reaction was run on a 1% formaldehyde 1.2% agarose gel in MOPS buffer (10X stock: 200mM MOPS, 50mM NaAc, 10mM Na_2_EDTA, pH 7.0) at 80V. Gels were stained with SYBR Gold (Invitrogen #S11494) for 30 minutes. For qRT-PCR and RNA-seq, the reaction was purified and size selected to > 200 nts using Zymo Clean and Concentrator-5 (Zymo #D4013) according to manufacturer’s protocol, eluting in 10μl of nuclease-free water. RNA was stored at -80°C.

### Poly(A) selection

Polyadenylated mRNA was selected using the NEBNext Poly(A) mRNA Magnetic Isolation Module (NEB #E7490L) according to the manufacturer’s protocol: 1 μg of total RNA was denatured at 65°C for 5 minutes then hybridized to buffered dT magnetic beads at room temperature for 2 minutes. Selected RNA was eluted in 50μl of Tris buffer at 80°C for 2 minutes and rehybridized to the same beads for a second round of selection at room temperature for 2 minutes. Re-selected RNA was eluted in a final volume of 17 μl of Tris buffer and stored at -80°C until further use.

### Quantitative reverse transcription PCR (qRT-PCR)

For first strand synthesis, a 20μl reaction consisting of Zymo-cleaned RNA (∼50 ng), dNTP (1uM), random primer (NEB #S1330S) (3uM), and DTT (10mM) was incubated at 65°C for 5 minutes then transferred to ice for 2 minutes. 1μl of SuperScript III reverse transcriptase enzyme and 8μl of 5X buffer (Invitrogen #18080085) were added to a final reaction volume of 40uL, incubated at 42°C for 90 minutes, then heat inactivated at 70°C for 15 minutes. Initial samples for the *X. laevis* 28S qRT-PCR were column purified (Qiagen #28704) and used at full concentration for qRT-PCR; subsequent samples were used directly at 1:10 dilution for qRT-PCR based on the results of a 4-sample, 1:5 dilution calibration curve analysis. qRT-PCR was performed in triplicate using 10μl reactions (2.5μl of cDNA, 5 μM of each forward and reverse primers, and 2x SYGreen mix (Genesee #17-505B)). qPCR was performed on QuantStudio 3 (Applied Biosystems) with an initial heat activation at 50°C for 2 minutes and then 95°C for 10 minutes. The reactions were cycled at 95°C for 15 seconds and 60°C for 1 minute for 40 cycles. Specificity was determined via a 3-stage melt curve analysis conducted at 95°C for 15 seconds, dropped to 60°C for 1 minute, and then raising the temperature from 60°C to 95°C at 0.1°C/s. No-template negative controls were run for each primer pair. Data analysis was conducted in Design and Analysis Application v1.5.1 (Thermo Fisher) and C_t_ values were calculated automatically from that application. Each NTC sample resulted in a C_t_ > 34. Experimental samples resulted in C_t_ values ranging between 14 – 33. ΔC_t_ values were calculated from the average of 3 technical replicates for each sample using mCherry as the reference gene and plotted ΔΔC_t_ values represent depletion conditions ΔC_t_ over input RNA control ΔC_t_. Statistical comparisons were done using two-tailed paired t tests on ΔC_t_ values (each treated sample is paired with the input RNA that was used for treatment). Primers were: **28S** (F – TGTGATTTCTGCCCAGTGCT ; R – GACGAGGCATTTGGCTACCT, amplicon: 107bp), **16S** (F – TCCAAAAACCTAGCATTCCAATTAT ; R – TTTCATCTTTCCTTACGGTACTTTTTC, amplicon: 140bp), **mCherry** (F – GCCCCGTAATGCAGAAGAAG ; R – TCAGCTTCAGCCTCTGCTTG, amplicon: 105bp), **sub1.L – XM_018266533.1** (F – AGCAGGAGAAATGAAGCCAGG – exon 4 ; R – CCGACATCTGCTCCTTCAGT – exon 5, amplicon: 80bp)[37] ; **helb.L – XM_018252426.1** (F – TTTCCAGGGTTCAGAAGAGGAG – exon12/13 junction ; R – TGCTATGGCTTCACCCAACT – exon 13, amplicon: 148bp) ; **nudt15.L – XM_018245539.1** (F – CCTGAGAAAAACGAAGGTTGGAA – exon3/4 junction ; R – TGGATTGTAGCCTTGCTGCT – exon 4, amplicon: 105bp). Primer specificity was verified using NCBI Primer-BLAST.

### RNA sequencing

Strand-specific RNA-seq libraries were constructed using the NEB Ultra II RNA-seq library kit (NEB #E7765) according to manufacturer’s protocol with fragmentation in first-strand buffer at 94°C for 15 minutes. Following first and second strand synthesis, DNA was purified with 1.8X AmpureXP beads (Beckman #A63880), end repaired, then ligated to sequencing adaptors diluted 1:5. Ligated DNA was purified with 0.9X AmpureXP beads and PCR amplified for 8 cycles, then purified again with 0.9X AmpureXP beads. Libraries were verified by Qubit dsDNA high sensitivity (Invitrogen #Q32851) and Fragment Analyzer prior to multiplexed sequencing (paired end 38/37bp) on an Illumina NextSeq 500 at the Health Sciences Sequencing Core at Children’s Hospital of Pittsburgh.

### RNA-seq data analysis

RNA-seq eads were mapped to the X. laevis v9.2 or GRCz11 (zebrafish) genomes using HISAT2 v2.0.5 [38] (--no-mixed --no-discordant) and assigned to genes (Xenbase v9.2 models for *X. laevis* and Ensembl r99 for zebrafish) using featureCounts v1.5.1 [39] in reversely-stranded paired-end mode with default parameters. To more accurately quantify rRNA levels in the *X. laevis* genome, due to poor assembly at the 40S rDNA locus, we additionally aligned to a separate HISAT2 index consisting of only the 40S (X02995.1) and 5S (J01009.1) sequences. Coverage plots were generated using BEDTools v2.25.0 genomeCoverageBed [40] and visualized on the UCSC Genome Browser[41]. To annotate histone mRNA, *X. laevis* and zebrafish protein sequences were curated from HistoneDB 2.0[42] and used to construct NCBI BLAST blastx databases[43]. Xenbase and Ensembl zebrafish mRNA hits with E-value < 1e-40 were annotated as histones. All plots and analyses were generated using R-3.4.4.

## RESULTS

### Sparse antisense oligo tiling effectively depletes rRNA

Previous RNaseH-based depletion methods used 50-nt DNA antisense oligomers that completely tile target RNA species[5, 24, 25] (Supplementary Fig S1a). We reasoned that for many applications, e.g. RNA-seq for non-degraded samples, tiling with gaps should be effective if the resulting fragments are short enough to be filtered out by size selection prior to cDNA generation. To test this strategy, we designed 39-40nt oligos spaced ≤ 30-nt apart (Fig 1a) to tile the *X. laevis* nuclear (28S, 18S, 5.8S, 5S) and mitochondrial (16S, 12S) rRNA. The ≤30nt gap ensures that a digested fragment would be <70nts long if the flanking oligos each induce cleavage in the center (Supplementary Fig S1b), smaller than most tRNAs. Shortening the oligo lengths allowed us to take advantage of value oligo pricing, and the overall strategy used 137 individual oligos tiling 5434 total bases (Supplementary Table S1), compared to 176 oligos and 8639 bases (a 37% reduction) for the 50-nt full-tiling strategy. This led to an 81% reduction in total oligo cost according to list prices (Methods). With the aid of a computational tool we created (see below), we ensured that most of the oligos had predicted melting temperatures (Tm) ≥65°C, with the exception of seven oligos targeting 16S rRNA with Tm between 58 and 64°C due to sequence constraints.

**Figure 1:**
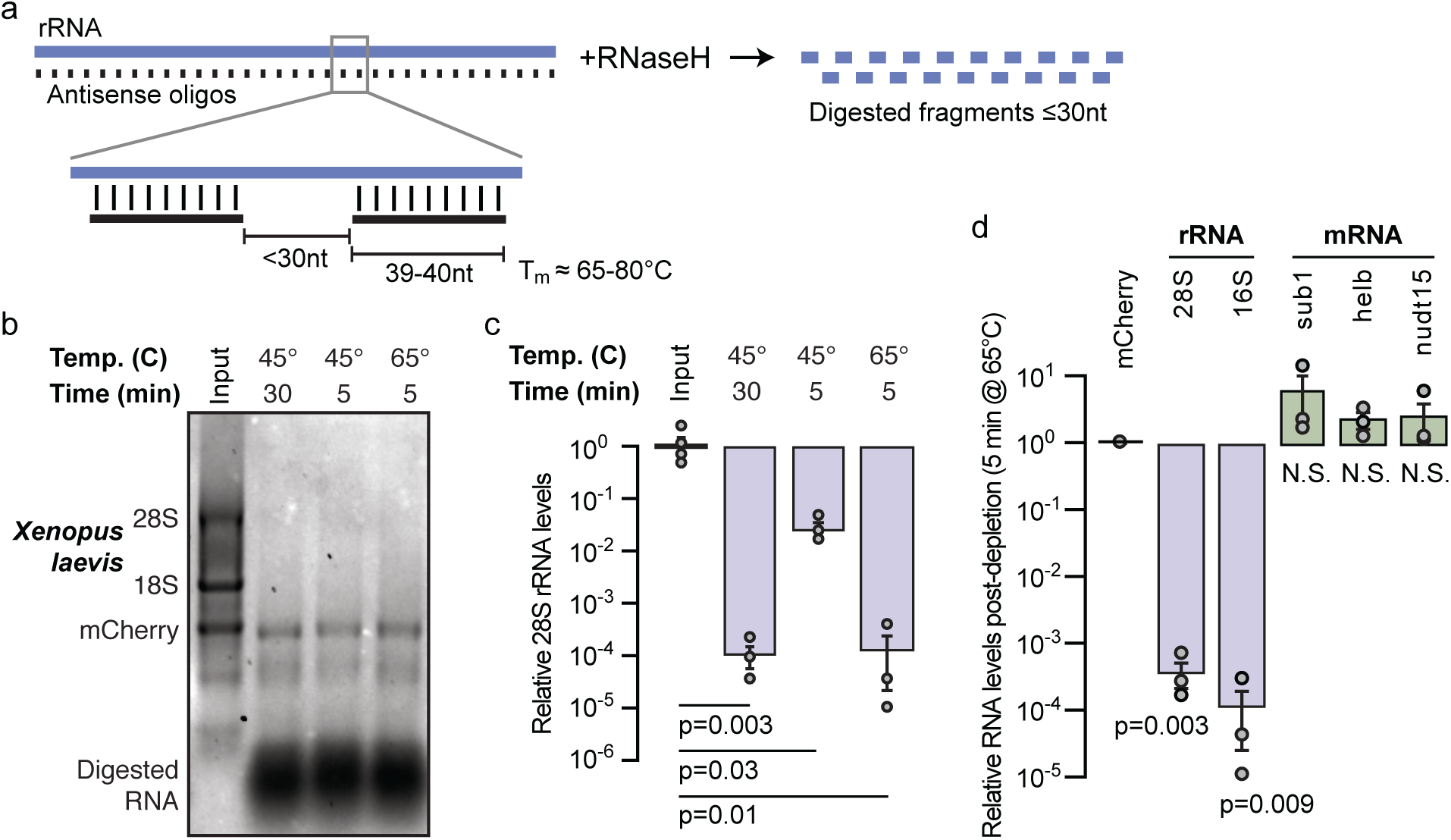
a) Schematic of rRNA depletion strategy using 39-40nt antisense oligos spaced ≤ 30-nt apart. b) *X. laevis* stage 0 total RNA (input, lane 1) and with rRNA depletion using different reaction conditions visualized on a 1% formaldehyde 1.2% agarose gel. In vitro transcribed mCherry mRNA was spiked into the input RNA prior to digestion. c) qRT-PCR comparing 28S rRNA levels in *X. laevis* stage 0 total RNA (input, left) versus depletion conditions normalized to mCherry. P values are from two-tailed paired t tests comparing depleted samples to their corresponding total RNA input. d) qRT-PCR measuring mCherry-normalized rRNA and mRNA levels in *X. laevis* stage 0 rRNA-depleted samples divided by levels in untreated samples. P values are from two-tailed paired t tests for each gene comparing depleted samples to their corresponding total RNA input. N.S. = not significant.

We combined aliquots of all of the nuclear rRNA-targeting oligos into a 10X working stock of 384μM, corresponding to 4 μM for each of the individual oligos; we created a similar stock for mitochondrial rRNA-targeting oligos at 1μM. At 1X, the oligo pools target ∼1 μg of total RNA, such that each oligo is in ∼10-fold excess of its rRNA target (Methods). To test the efficacy of the oligo pools, we subjected Nieuwkoop and Faber (NF) embryonic stage 0 *X. laevis* total RNA to RNaseH treatment, followed by Turbo DNase, and visualized the digested RNA without any cleanup on a 1% formaldehyde-agarose gel. We tested previously published reaction parameters (45°C for 30 minutes) using New England Biolabs thermostable RNaseH along with two other conditions that reduced reaction time (45°C for 5 minutes) and additionally increased reaction temperature (65°C for 5 minutes) (Fig 1b). To test specificity of the treatment for rRNA, we spiked 150 ug of in vitro transcribed mCherry mRNA into each reaction. All three reaction conditions were effective, eliminating the upper bands corresponding to the 28S (4082 nts) and 18S rRNA (1825 nts) while leaving the mCherry (1037 nts) band intact (Fig 1b). A diffuse band migrating at ∼500nts is also intact in the digested samples, which likely corresponds to highly abundant histone mRNA species based on inspection of RNA-seq datasets. A large mass that is likely digested RNA and DNA oligos is visible at the bottom of each lane at <50nts (Fig 1b), which we expect to be largely excluded if size selection is performed after digestion.

To precisely quantify the rRNA depletion, we subjected samples in triplicate to quantitative reverse transcription PCR (qRT-PCR) probing for 28S rRNA. All three depletion conditions significantly reduce the level of 28S rRNA compared to untreated RNA, with the 45°C/30 minute and 65°C/5 minute reactions reducing 28S rRNA levels by 99.99% (p ≤ 0.01, two-tailed paired t test) (Fig 1c) – the optimal reaction temperature for the thermostable RNaseH is 65°C, and these results demonstrate that digestion is rapid at this temperature.

To assess the effects of rRNA depletion on mRNA as compared to rRNA, we performed qRT-PCR on treated (RNaseH 65°C/5 minutes) versus untreated total RNA, probing for embryonic mRNA expressed at low to moderate levels based on previous RNA-seq studies[44] – *sub1.L* (133 transcripts per million (TPM)), *helb.L* (5 TPM), and *nudt15.L* (1 TPM) – along with 28S and mitochondrial-encoded 16S rRNA, normalizing to mCherry spike in. Both rRNA species were significantly depleted in treated versus untreated samples (p < 0.001, two-tailed paired t test) (Fig 1d), while the mRNA levels were not significantly different (p > 0.1, two-tailed paired t test) (Fig 1d). Taken together, we find that the optimized oligo design effectively and specifically degrades targeted rRNA, using streamlined reaction times.

### Optimized rRNA depletion yields high quality RNA-seq libraries

Next, we sought to determine whether our depletion strategy could be used to construct high quality RNA-seq libraries. We collected total RNA from two different *X. laevis* embryonic stages (NF 5 and 8) and performed either rRNA depletion (65°C/5 minutes) or poly(A)+ selection, then built Illumina strand-specific libraries and sequenced each sample to 10 million read pairs. Both the poly(A)+ and rRNA depleted samples show a >100-fold reduction in reads aligning the 40S rDNA locus compared to unselected total RNA (Fig 2a). Indeed, >90% of reads align to annotated mRNA or long non-coding RNA (lncRNA) in both the poly(A)+ and rRNA depleted samples, compared to <5% for unselected total RNA (Fig 2b, Supplementary Fig S2a).

**Figure 2:**
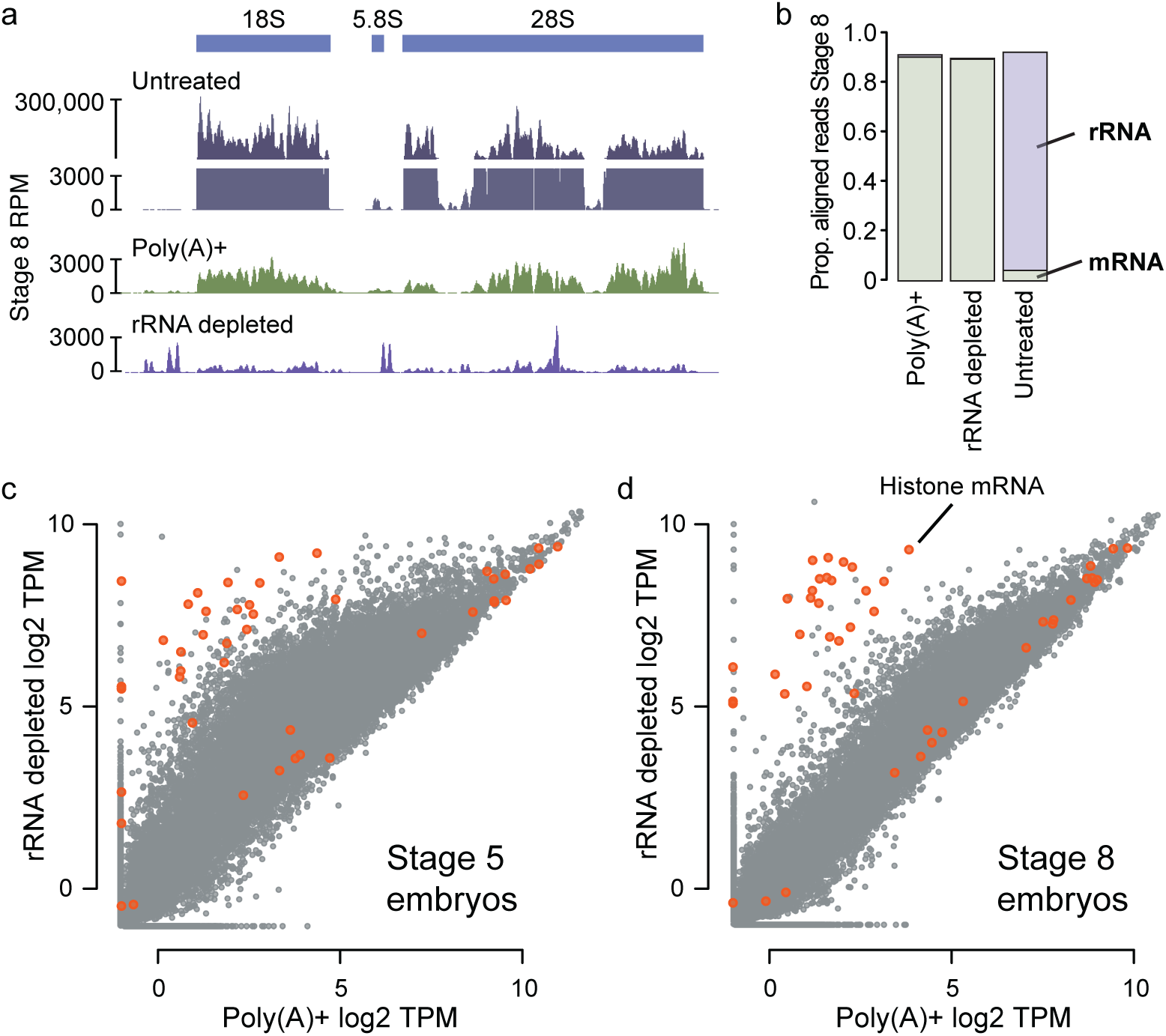
a) Genome browser tracks comparing read coverage at the *X. laevis* 40S rDNA locus in untreated total RNA, poly(A)+ and rRNA depleted RNA-seq libraries from stage 8 embryos. Y-axis is discontinuous for the total RNA sample. b) Stacked barplots showing proportion of aligned reads deriving from mRNA (green) versus rRNA (purple) in untreated, poly(A)+ and rRNA depleted RNA-seq libraries. c,d) Biplots comparing log2 TPM expression levels from poly(A)+ and rRNA-depleted libraries at stage 5 and 8, respectively. Histone genes are highlighted in orange.

Transcriptome wide, expression levels correlate well between poly(A)+ and rRNA depletion for most genes (Fig 2c,d, Supplementary Table S3). However, at stage 5 a population of transcripts shows elevated apparent levels with rRNA depletion compared to poly(A)+ (Fig 2c). Indeed, the maternal RNA contribution to the egg is largely deadenylated, with poly(A) tails lengthening during early embryonic stages through cytoplasmic polyadenylation[11]. Thus, rRNA depletion avoids the depressed expression levels arising from inefficient capture of mRNA with short poly(A) tails, typical of poly(A)+ RNA-seq[12-15]. By the mid-blastula transition (NF stage 8), poly(A) tails are longer, so poly(A)+ and rRNA depletion yield comparable expression values for these mRNA (Fig 2d). However, some RNA species are still better represented in the rRNA depletion libraries, suggesting these transcripts lack poly(A) tails. Indeed, replication-dependent histone mRNA encode 3’ stem loops instead of poly(A) tails[45], and we find these transcripts are much more efficiently sequenced with rRNA depletion (Fig 2c,d). Thus, our optimized rRNA depletion strategy effectively quantifies expression levels of both the adenylated and non-adenylated transcriptome.

### Compact oligo pools can simultaneously target divergent rRNAs

Given the gapped design strategy, it is likely that some sequence differences in target RNAs would be tolerated, allowing oligo pools designed for the rRNAs of one taxon to be used for another closely related taxon. At greater sequence dissimilarity, we reasoned that shared oligos could be designed to target common subsequences between two or more RNAs, with gaps positioned over variable regions, avoiding the need to design completely separate reagents for rRNA depletion.

To test this, we designed a combined oligo pool to target the two versions of the zebrafish nuclear rRNAs, which are 86% similar. Zebrafish encode maternal-specific 28S, 18S, 5.8S and 5S rRNAs that are deposited into eggs during oogenesis[35, 36]. After zygotic genome activation, distinct somatic rRNAs begin to be transcribed and slowly replace the maternal versions as the embryo develops (Fig 3a). Thus, to effectively deplete rRNAs in zebrafish embryos, both versions would need to be targeted. We aligned each rRNA sequence pair and designed 46 oligos that target identical regions between the maternal and somatic versions. To target regions that differed at only one position, we additionally designed 22 oligos containing a wildcard base (e.g., R to represent either A or G), which are ordered as mixtures of two oligos (Fig 3b). Finally, 44 and 41 additional oligos were required to target divergent maternal and somatic regions, respectively. In all, the combined design required 153 total oligos to together target both sets of nuclear rRNAs (Supplementary Table S2), compared to 201 total oligos for two independent sets.

**Figure 3:**
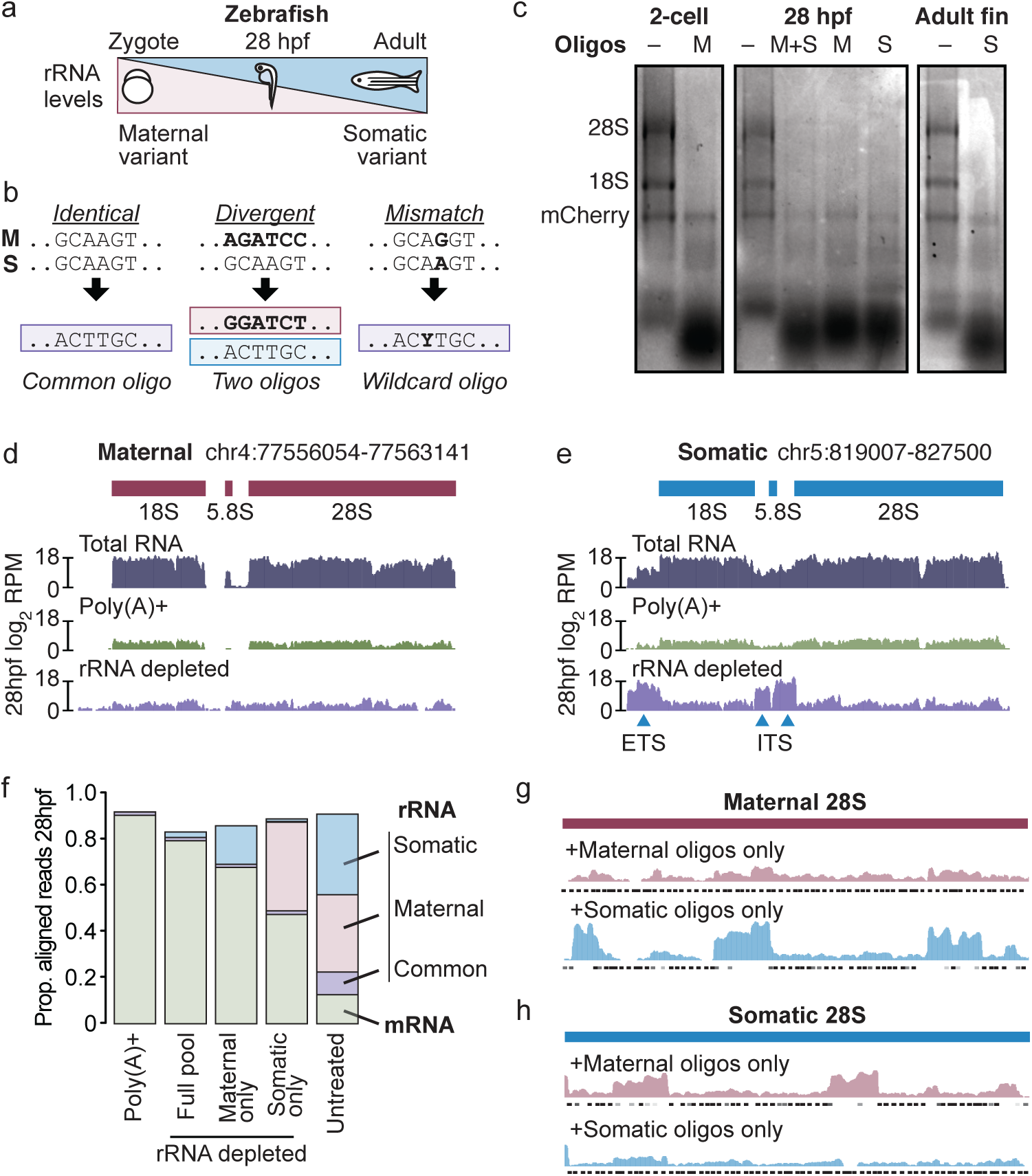
a) Diagram illustrating the relative expression of the maternal and somatic nuclear rRNA variants over development. hpf = hours post fertilization. b) Schematic showing how oligos (bottom) can target similar sequences (top) between two RNAs. c) Gels showing total RNA samples after rRNA depletion across three zebrafish developmental timepoints using only the maternal pool (M), only the somatic pool (S), or a mixture of the two pools (M+S), as compared to untreated input (-). In vitro transcribed mCherry mRNA was spiked into the input RNA prior to digestion. d,e) Genome browser tracks comparing read coverage at the maternal (d) and somatic (e) 45S rDNA loci in untreated, poly(A)+ and rRNA depleted libraries from zebrafish 28hpf. f) Stacked barplots showing proportion of aligned reads deriving from mRNA (green) versus rRNA (blue, uniquely somatic; pink, uniquely maternal; purple, common) in untreated, poly(A)+ and rRNA depleted RNA-seq libraries. g & h) Genome browser tracks comparing read coverage at the maternal (g) and somatic (h) 28S rDNA loci in rRNA depletion libraries depleted using only maternal (top row) or only somatic (bottom row) oligo pools from zebrafish 28hpf. Targeted regions by each oligo pool are shown beneath each track.

We combined the common and unique oligos to create separate maternal and somatic pools each at 4μM per individual oligo. We also created a mitochondrial rRNA-targeting pool at 1μM per oligo (there is only one known version of the 16S and 12S rRNAs). In the 2-cell stage embryo, the maternal+mitochondrial pools effectively and specifically induce rRNA depletion from total RNA, which is entirely maternally derived (Fig 3c, left), while in adult fins, the somatic+mitochondrial pools are effective (Fig 3c, right). We additionally tested depletion in 28 hours post fertilization (hpf) embryos, which express roughly equal amounts of maternal and somatic rRNA [36]. Neither the maternal pool nor the somatic pool alone was as effective as a 1:1 mixture of both pools: using only the maternal or somatic pools produced several RNA species between 300 and 800 nts, suggesting incomplete digestion (Fig 3c, middle).

To quantify this difference in efficiency, we constructed RNA-seq libraries at 28hpf. rRNA depletion with the combined oligo pool effectively reduced the number of sequencing reads mapping to either the maternal or somatic rRNA loci compared to untreated total RNA, comparable to poly(A)+ (Fig 3d,e, Supplementary Fig S2b). We did observe elevated levels of reads mapping to the external and internal transcribed spacers of the full somatic 45S transcript (5’ ETS and two ITS regions; Fig 3e), which were omitted from the oligo design; as well as a small region of 16S rRNA where targeting was less efficient (Supplementary Fig S2c-e). Nonetheless, 79% of reads mapped to mRNA or lncRNA, compared to 90% for poly(A)+ (Fig 3f), and expression quantification was highly correlated between the two methods (Supplementary Fig S2f, Supplementary Table S4). rRNA depletion additionally recovered highly expressed non-coding RNAs such as the signal recognition particle and 7SK RNAs, which are not efficiently sequenced with poly(A)+ (Supplementary Fig S2f).

In contrast, rRNA depletion using only the maternal or somatic pools was less efficient. By targeting only the maternal rRNA, 15% of the library is still rRNA, mapping to the somatic 45S locus; and by targeting only the somatic rRNA, 38% of reads derive from rRNA, corresponding to the maternal 45S locus. This leaves only 68% and 47% of the library mapping to mRNA+lncRNA, respectively (Fig 3f). Read coverage over the maternal 28S rRNA gene indeed shows a failure to digest sequence regions where the somatic oligos lack complementarity (Fig 3g), while a similar pattern is observed over the somatic 28S gene when only maternal oligos are used (Fig 3h). These results show that a full maternal+somatic targeting strategy is required to achieve a maximally effective rRNA depletion and demonstrate that a shared, compact oligo pool can efficiently target these divergent sequences simultaneously.

### The Oligo-ASST Web tool streamlines antisense oligo design

The lack of an appropriate computational method to implement a gapped tiling strategy prompted us to build a Web tool called Oligo-ASST (which stands for <u>A</u>nti<u>s</u>ense <u>S</u>paced <u>T</u>iling) using the Python Dash v1.0 framework, available at https://mtleelab.pitt.edu/oligo. Oligo-ASST iteratively positions antisense oligos along a target sequence to maximize distance between consecutive oligos up to a threshold (e.g., 30 nts) while attempting to maintain a predicted Tm as close to 70°-80°C as possible according to RNA/DNA duplex thermodynamic parameters[33]. To design oligos, the user uploads one or more sequences in FASTA format (Fig 4a), selects oligo and gap length parameters according to their needs, then the resulting oligo sequences, coordinates and properties are displayed in the Web interface, where they can be downloaded in text format (Fig 4b, right, Supplementary Fig S3). Tiled positions are also highlighted on a Dash Bio Sequence Viewer (Fig 4b, left).

**Figure 4:**
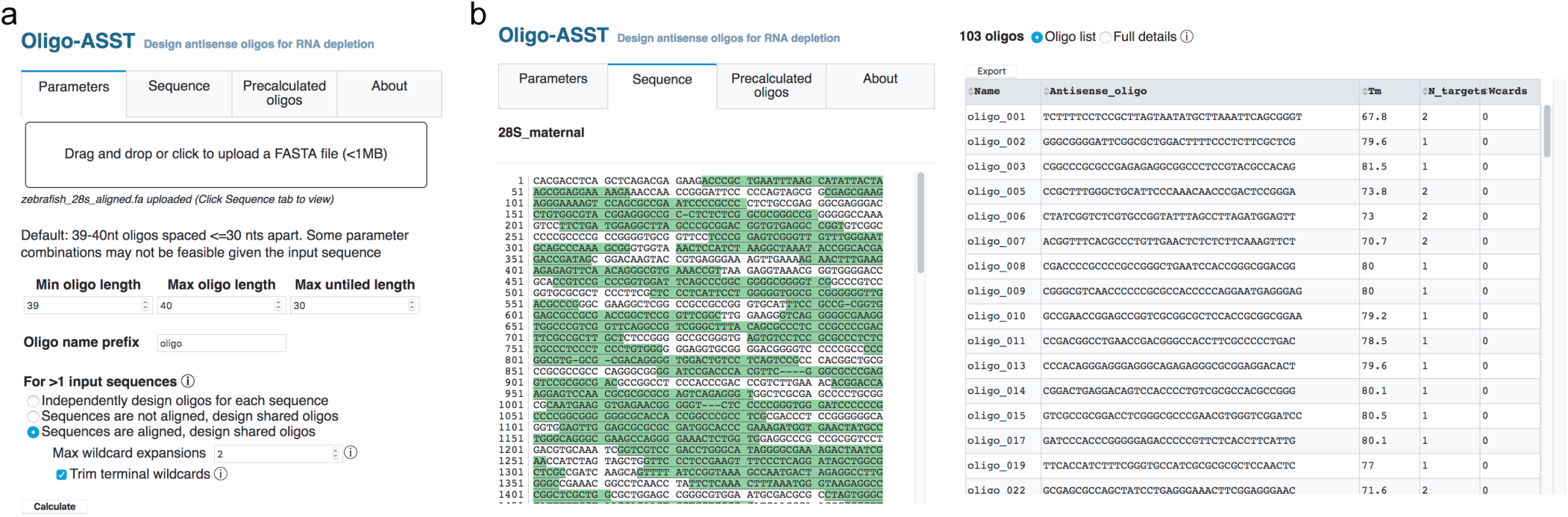
a) The Oligo-ASST Web interface allows users to upload a FASTA file for target sequences and select parameters for oligo design. b) Designed oligos are highlighted in a sequence viewer (left) and listed in the right pane in abbreviated form or with full details (not shown), which can be downloaded in text format.

When multiple sequences are input, users can choose to design independent oligos per sequence or a shared set with common oligos targeting either identical subsequences or subsequences with one or more mismatches using wildcard bases. Sequences can be aligned beforehand using a tool such as MUSCLE[46] to improve identification of identical subsequences and yield a maximally compact oligo pool to target heterogeneous RNA species.

## DISCUSSION

Here, we demonstrate that a streamlined RNaseH digestion protocol using easily obtained reagents efficiently and cost-effectively achieves ribosomal RNA depletion, which we estimate to be ∼$8 per reaction. Our Web tool Oligo-ASST improves oligo design to use shorter antisense DNA oligos (39-40 nts) that tile rRNA target sequences with gaps, thereby reducing reagent cost compared to previous methods [5, 24] while still producing high-quality RNA-seq libraries comparable to those constructed with poly(A) selection (Fig 2-3). Although there may be use cases where magnetic bead-based methods would be more appropriate, e.g. highly degraded RNA[6] or libraries where precise ends are required such as for ribosome profiling[47], for many RNA-seq applications RNaseH digestion should yield excellent results. In addition, since Oligo-ASST can also design compact oligo sets for treating multiple different rRNAs by targeting shared sequences, this protocol is especially advantageous for researchers to achieve rRNA depletion in diverse taxa.

We found that gaps of ≤30nts untargeted by oligos did not seem to affect overall performance of rRNA depletion, suggesting that the majority of the resulting digested fragments are too short to be retained after column cleanup and size selection for the RNA-seq libraries. Thus, we have demonstrated that the previously standard full tiling strategy is unnecessary for a typical RNA-seq use case. Increasing allowable gap length is likely to leave larger digested fragments and reduce efficiency of the depletion (Supplementary Fig S2c,d), though there may be applications for which this may be acceptable. Conversely, decreasing allowable gap length could facilitate recovery of smaller RNAs by ensuring that digested fragments are more easily size separable from the desired RNA species.

Our digestion reaction proceeds for only 5 minutes at 65°C using NEB thermostable RNaseH. Reactions at lower temperatures seem to produce comparable results, which will be beneficial when targeting AT-rich RNAs with lower Tms. However, it is likely that the higher reaction temperature reduced the likelihood of off-targeting by the oligos. Indeed, we found no evidence in our RNA-seq libraries that non-rRNA gene quantification was affected due to treatment (Fig 2d, Supplementary Fig S2), additionally justifying our use of shorter targeting oligos. Future optimizations to oligo design could avoid targeting regions with high sequence similarity to non-rRNAs, which is possible due to the flexibility of the gapped oligo tiling strategy.

For many taxa, designing new oligo sets should be straightforward with Oligo-ASST, given the availability of rRNA sequences in databases such as GenBank. For *Xenopus* and zebrafish, we found that the majority (75-86%) of rRNA reads in total RNA derive from 28S and 18S rRNA (Supplementary Fig. S2a,b), thus targeting these alone would still yield RNA-seq libraries with a majority of non rRNA-reads. However, we did find a substantial fraction of reads mapping to 5.8S rRNA as well as the transcribed spacer regions ITS1, ITS2, and the 5’ ETS of the pre-rRNA; and the 16S and 12S mitochondrial rRNA, indicating that a maximally comprehensive oligo pool would target at least each of these eight sequences. Depending on the transcriptome of interest, it may also be valuable to target other abundant RNAs for depletion, e.g. the 7SK small nuclear RNA (Supplementary Fig S2f); inspection of existing RNA-seq libraries would reveal such RNA species. Indeed, Oligo-ASST is agnostic to RNA identity and can be used to design oligos that target arbitrary sequences.

It is likely that targeting reagents will be somewhat robust to polymorphisms in the rRNA sequences, which may be especially prevalent in non in-bred strains and species. Indeed, we demonstrate that Tms as low as 58°C still seem to be effective, suggesting that a small number of mismatches could be tolerated. However, Oligo-ASST can facilitate the design of oligos that map to non-variable regions, which as we show in zebrafish, allows a partially overlapping oligo pool to simultaneously target the two divergent sets of rRNAs in the zebrafish genome.

In conclusion, we have developed and optimized antisense oligo-based rRNA depletion for *X. laevis* and zebrafish RNA-seq libraries and provide a tool Oligo-ASST to design similar reagents for any other species. We anticipate this will be of benefit to researchers who need alternatives to poly(A) selection for RNA-seq, particularly those working with taxa that were inadequately served by previous rRNA depletion methods.

## AVAILABILITY

The Oligo-ASST Web tool is available at https://mtleelab.pitt.edu/oligo.

Source code for the Web application and a command-line version of the program are available at https://github.com/MTLeeLab/oligo-asst.

## ACCESSION NUMBERS

All raw sequencing reads are deposited in the NCBI Gene Expression Omnibus under accession number GSE152902.

## SUPPLEMENTARY DATA

Supplementary Figures S1-3

Supplementary Table S1: Antisense oligos for *X. laevis* rRNA depletion

Supplementary Table S2: Antisense oligos for zebrafish for rRNA depletion

Supplementary Table S3: Transcripts per million expression levels for *X. laevis* RNA-seq experiments

Supplementary Table S4: Transcripts per million expression levels for zebrafish RNA-seq experiments

## AUTHOR CONTRIBUTIONS

WAP & MTL conceived of the project, performed the experiments, and wrote the manuscript. MTL wrote the software. AEC oversaw *Xenopus* embryo collection.

## ACKNOWLEDGEMENTS

We thank J. Rosenbaum, T. Ayers, L. Wolf, R. Bainbridge, M. Ellison, S. Hainer, B. Moore, P. Rangan, and the entire Lee lab for technical assistance and discussions. This project used the University of Pittsburgh Health Sciences Sequencing Core at UPMC Children’s Hospital of Pittsburgh for sequencing.

## FIGURE LEGENDS

**Supplementary Figure S1:**
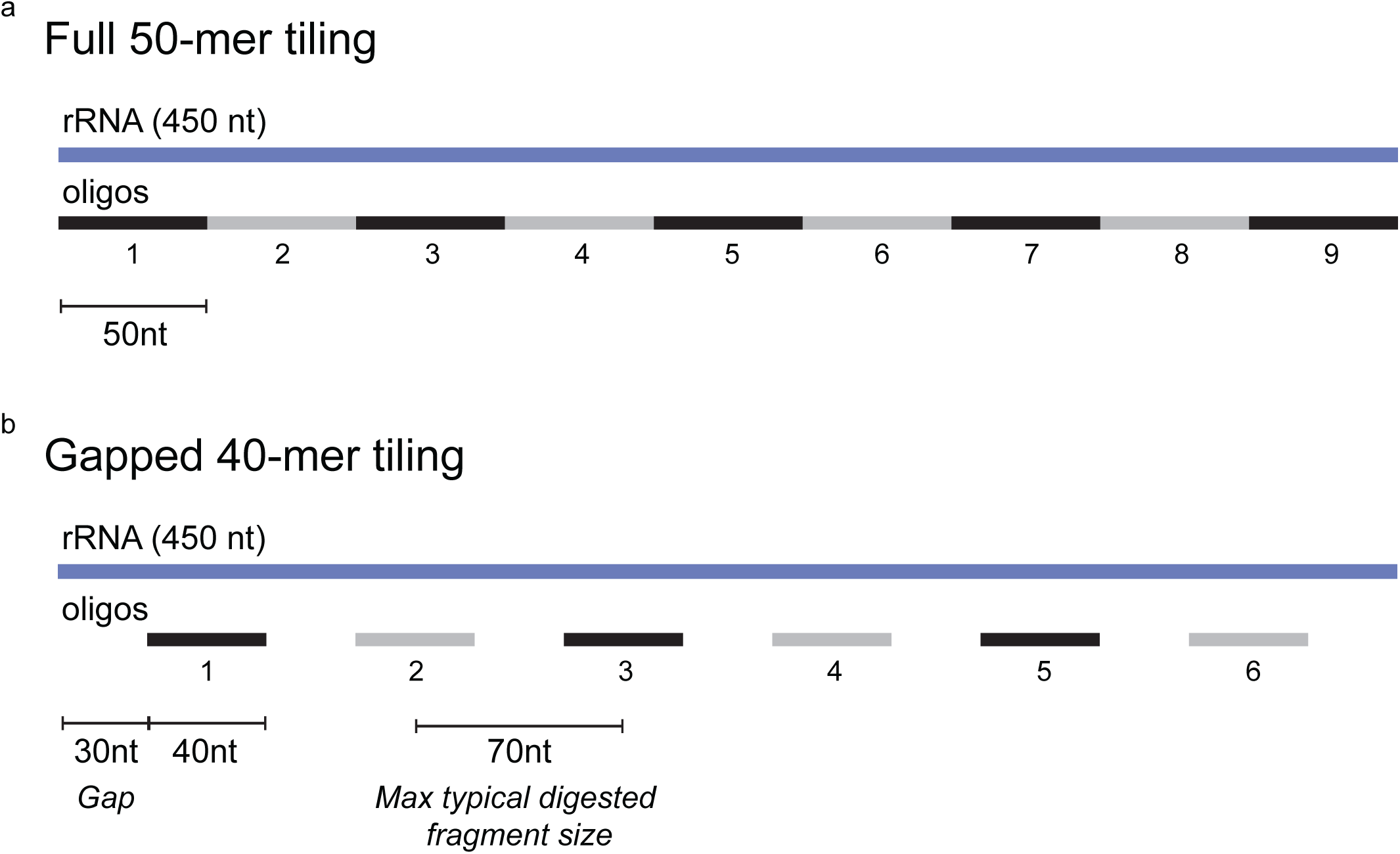
a) Traditional design for rRNA depletion using 50mer antisense oligos that fully tile the target rRNA. b) A gapped tiling design using 40mer antisense oligos with 30nt gaps uses fewer oligos to digest the target rRNA to <70nt.

**Supplementary Figure S2:**
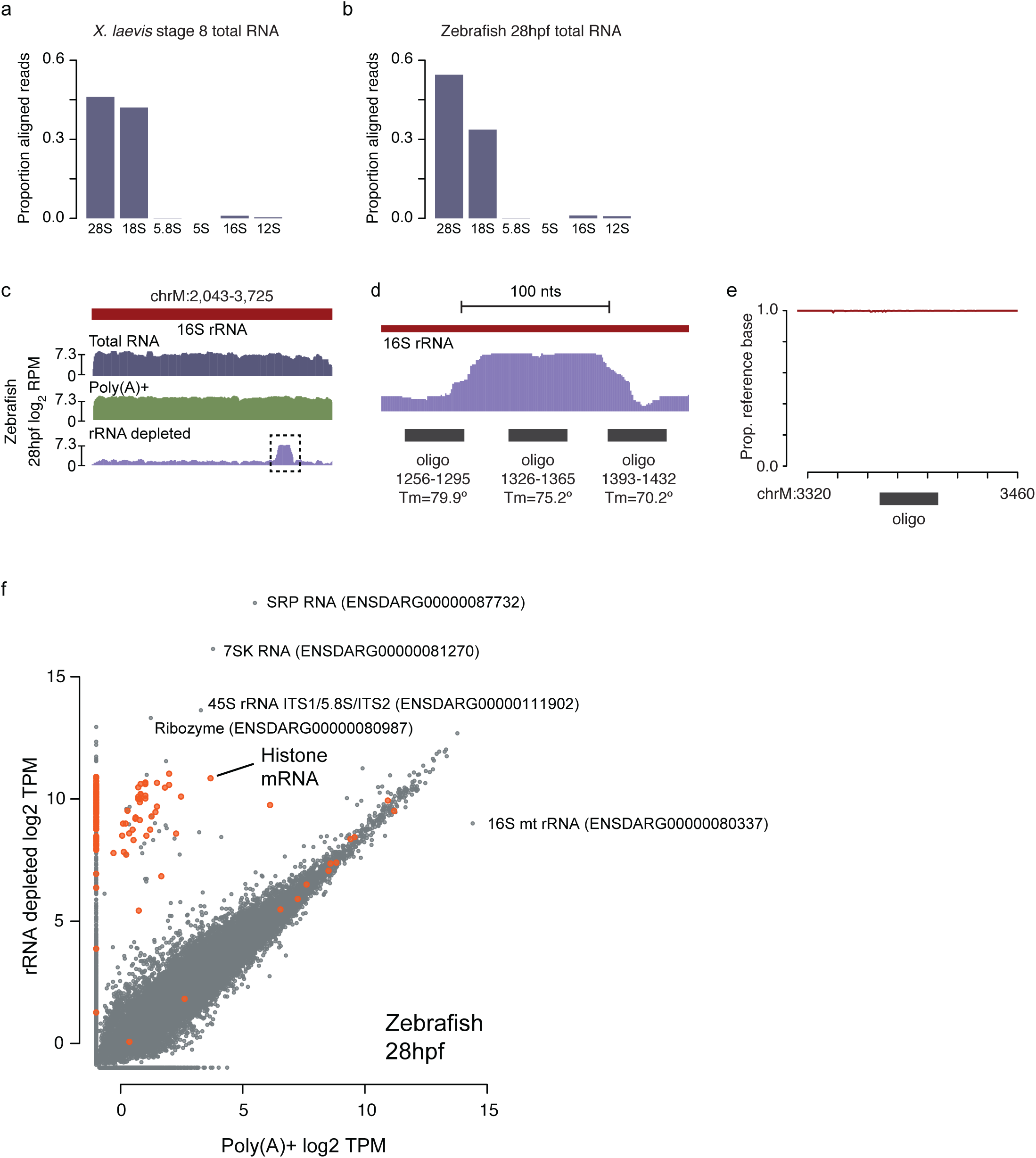
a,b) Barplots showing proportion of aligned reads from untreated total RNA mapping to rRNA species in *X. laevis* (a) and zebrafish (b). c) Genome browser tracks comparing read coverage at the 16S rDNA loci in untreated, poly(A)+ and rRNA depleted libraries from zebrafish 28hpf embryos. One region with less efficient rRNA depletion is boxed. d) Zoomed browser track for the boxed region in (b) showing that the inefficient digestion occurred over the region targeted by one oligo (positions 1326-1365 relative to the 16S sequence). This likely resulted in rRNA fragments that were slightly too large to be efficiently excluded during cleanup and library build. e) Variant analysis of sequencing reads mapping to chrM:3320-2460 showing that nearly all read sequences match the GRCz11 reference sequence, suggesting that there is no defect in the oligo’s ability to target; rather, it is likely that this oligo was omitted from the pool in error. f) Biplot comparing log2 TPM expression levels from poly(A)+ and rRNA-depleted libraries at 28hpf. Histone genes are highlighted in orange. Several highly expressed non-coding RNAs are labeled.

**Supplementary Figure S3:**
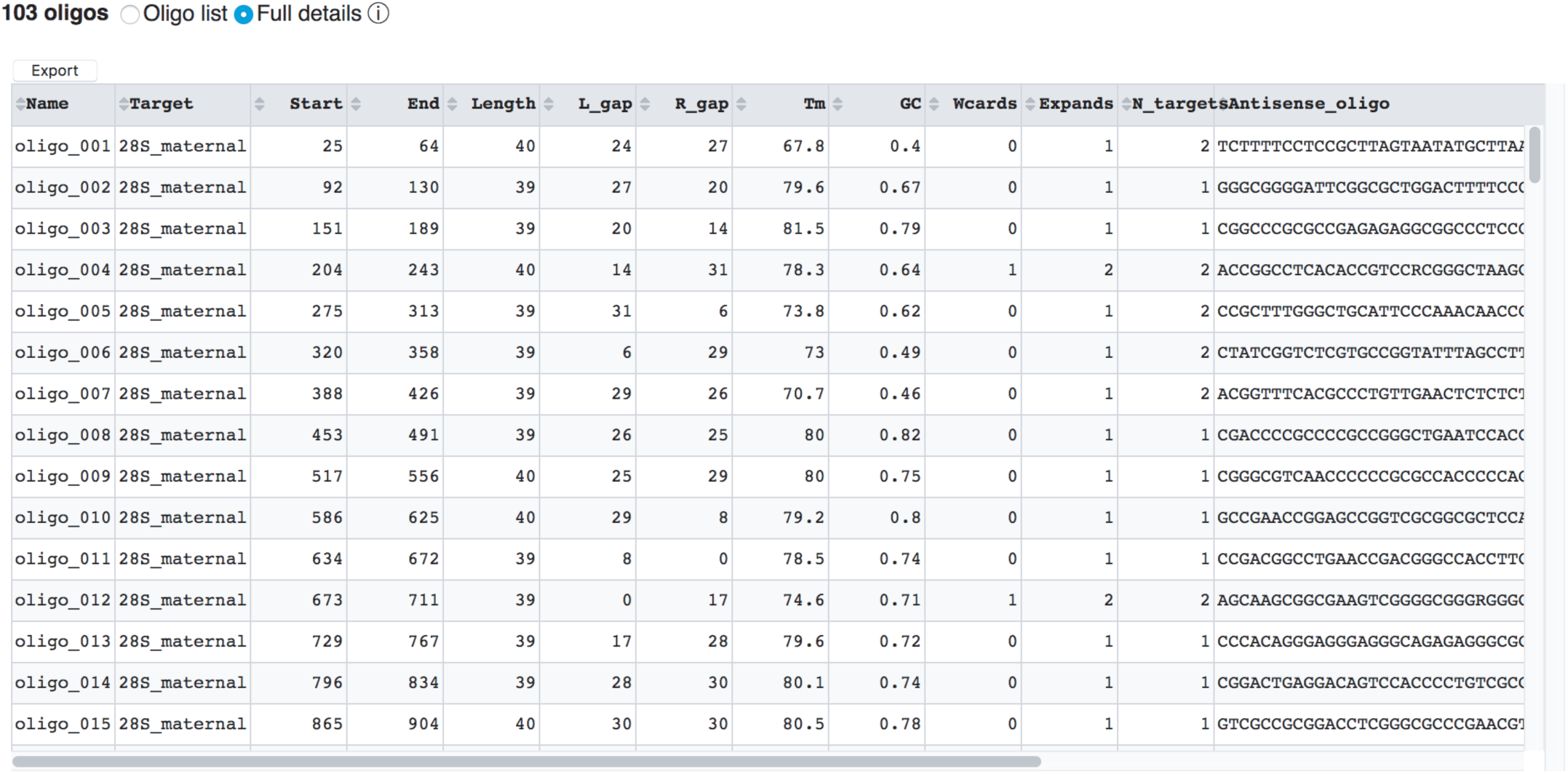
Oligo-ASST Web screenshot of the detailed results from designing antisense oligos targeting the zebrafish maternal and somatic 28S rRNAs together.

